# A Latent Inflammatory Tissue-State Variable Mechanistically Links Radiotherapy-Induced Immune Remodeling to Recurrent Tumor Permissiveness

**DOI:** 10.64898/2026.09.14.751444

**Authors:** McKenzie A. Mayeaux, Xin Maizie Zhou, Marjan Rafat

## Abstract

Triple-negative breast cancer recurrence following radiotherapy is associated with a microenvironment characterized by immune dysfunction and persistent inflammation. We developed an experimentally constrained agent-based model to investigate how transient immune remodeling becomes a persistent recurrence-permissive tissue state. The model reproduced experimentally observed macrophage recruitment and phenotype dynamics but demonstrated that recurrent recruitment, impaired inflammatory resolution, and adaptive immune bias were insufficient to reproduce the recurrent macrophage ecology. We therefore introduced recurrence-associated microenvironmental inflammation (RAMI), a latent tissue-state variable representing accumulated unresolved inflammatory remodeling. Coupling RAMI to the emergence of experimentally constrained interleukin-6 signaling generated tissue-to-cell feedback that reinforced recurrence-associated macrophage phenotypes and increased tumor establishment, supporting inflammatory tissue memory as a mechanistic intermediary between transient immune perturbation and persistent recurrent tumor permissiveness.

## Introduction

Inflammation following tissue damage is necessary to clear unwanted cell debris, bacteria, and other foreign particles that are introduced during injury. It is expected that immune cells, such as macrophages, neutrophils, and T cells, that initially flood the site with inflammatory factors will abate. The wound typically exits acute inflammation, giving way to wound healing immune cell phenotypes which calm the inflammatory reaction and secrete factors and extracellular matrix (ECM) components. Eventually, this leads to resolution with tissue repair^1–3^. However, some tissues do not resolve and instead continue to experience the effects of lingering immune cells and aberrant inflammation called chronic inflammation. For these tissues, wound healing and resolution are stalled, leading to pain, loss of function, and other deleterious impacts on patient health.

Patients with triple-negative breast cancer (TNBC) recur at higher rates and experience worse outcomes relative to other breast cancer subtypes^4^. Clinical data has identified immunodeficiency following radiotherapy to be associated with TNBC recurrence, characterizing a particularly vulnerable patient population^5^. Here, macrophages have been identified as the central drivers of recurrence, and their increased recruitment and M2-like phenotype was found to increase tumor cell infiltration into the irradiated site^5,6^ (**Figure 1A**). Experimental evidence presents a seemingly counterintuitive microenvironment where immunodeficiency and chronic inflammation after radiotherapy cooperate to promote a recurrence-permissive milieu^6–8^ (**Figure 1B**).

**Figure 1.**
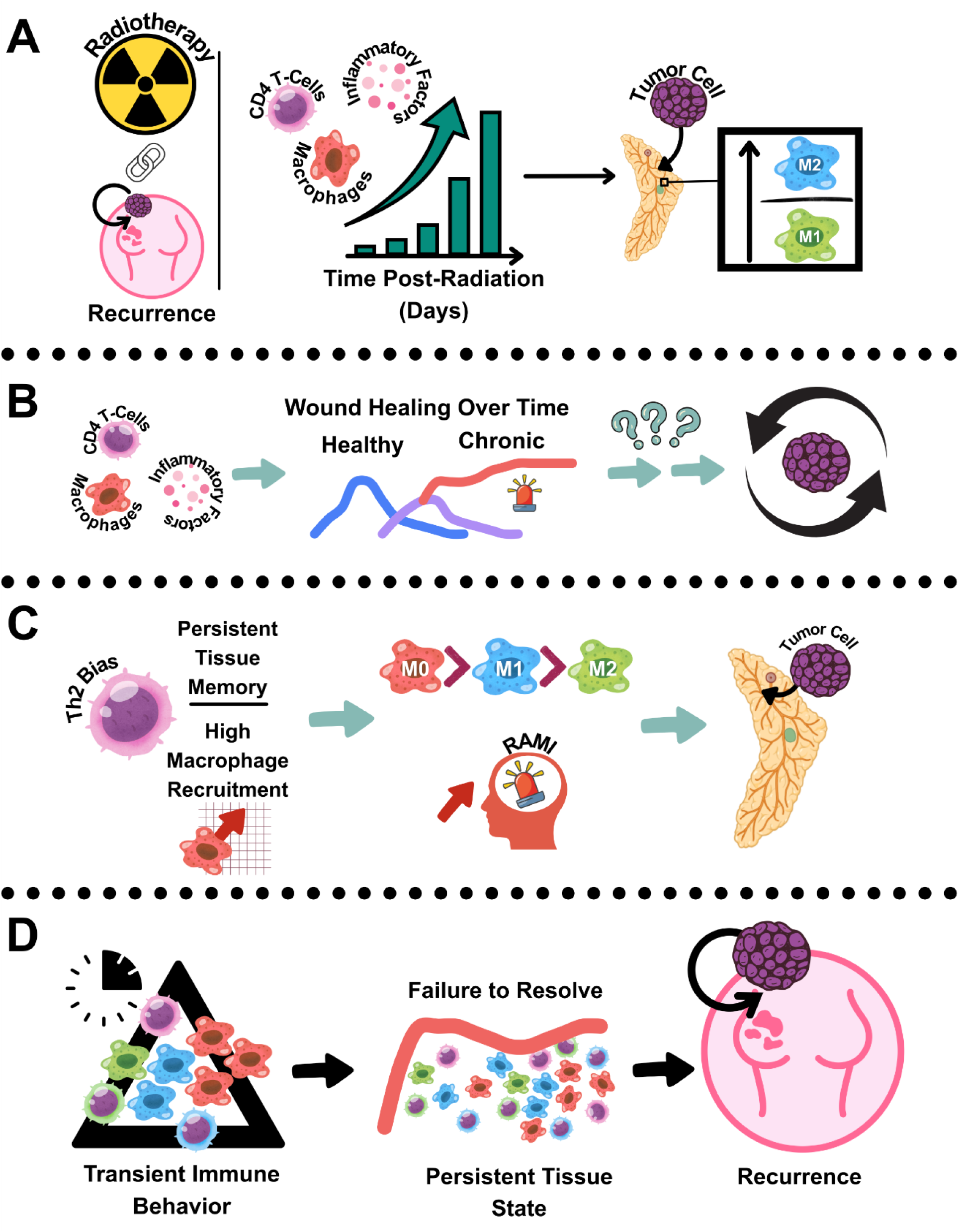
Biological motivation for modeling inflammatory tissue memory following radiotherapy. Radiotherapy can create a paradoxical tissue state in which chronic inflammation and immune dysfunction coexist to promote recurrence. (A) Experimental observations motivating the model. (B) Knowledge gap. (C) Computational hypothesis. (D) Proposed mechanism linking transient immune remodeling to persistent tissue-state changes.

**Figure 2.**
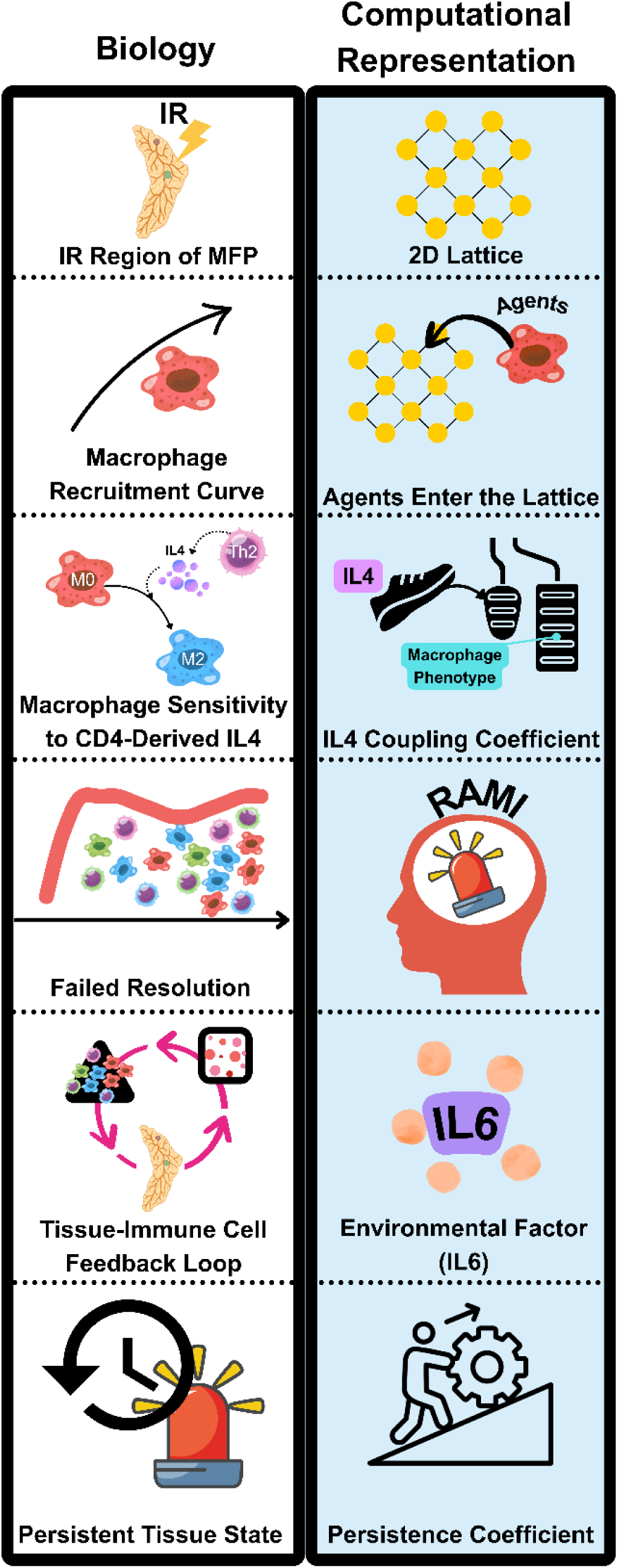
Computational representation of the recurrent tissue microenvironment. Biological components of the irradiated tissue were translated into computational abstractions, including macrophages as agents, irradiated tissue as a lattice, inflammatory tissue remodeling as the latent state variable recurrence-associated microenvironmental inflammation (RAMI), interleukin-6 (IL6) as an emergent inflammatory mediator, and tumor permissiveness as a latent tissue property.

Radiotherapy, though a powerful therapeutic tool, can inflect damage to normal tissue surrounding the treatment area. It has been established that systemic inflammation for such patients causes an increase in the occurrence of progression, recurrence, and patient mortality^9^. Given that radiotherapy is a key therapy that most patients will receive as a part of their treatment regimen, it is important to understand why immunodeficient TNBC patients experience prolonged inflammation and failed resolution post-therapy^5^.

While molecular approaches have identified many of the granular tissue and cellular behaviors that characterize this environment, it remains challenging to understand how transient immune perturbation following radiotherapy in the context of immunodeficiency contributes to persistent inflammation and recurrence^6–8^. Macrophages, especially those with an M2-like phenotype, have been given much consideration in cancer therapies due to their known role in wound healing following radiotherapy^10^, their plasticity^11^, targetability in wound healing^12^,and association with poor outcomes^5,6,13^. Stil, a systems biology examination of the site would yield crucial information about how the tissue switches from a nonrecurrent to recurrent state^14,15^.

To address this, we have developed an experimentally grounded agent-based model (ABM) that represents the irradiated tissue as a dynamic system rather than a collection of isolated cell behaviors^14,16^. These experimental observations constrained macrophage recruitment^5^, phenotype dynamics^6^, and CD4+ T-cell composition^17^, providing quantifiable biological anchors for hypothesis-driven computational experiments investigating immune bias, tissue memory, inflammatory persistence, and recurrence (**Figure 1C**).

Many ABMs tend to focus on direct, cell-cell interactions or immediate mechanistic outcomes^18^. However, computational models increasingly use mechanistic abstractions to represent biologically meaningful processes that are not directly observable^19^. Our model simulates how cells lead to an altered tissue state that impacts future cell behavior by introducing a latent tissue-state variable that captures the cumulative consequences of immune perturbation and inflammatory remodeling within the irradiated microenvironment. This is particularly helpful for this recurrent niche because it is the persistence of chronically inflamed tissue that seems to be central to recurrence after radiotherapy. Modeling inflammation and wound healing is well-established^14,19,20^. Here, we use a latent tissue-state variable to integrate inflammatory events into tissue outcomes. Rather than solely modeling direct interactions among cells, the model represents how immune cells and the environment alter each other to sustain inflammation and ultimately permit tumor cell colonization of tissues (**Figure 1D**).

## Methods

### Model Overview

The model represents a finite 2D cross-section of irradiated mammary tissue corresponding to the tissue surrounding the treatment field during radiotherapy for TNBC. The bounded lattice reflects the finite region of tissue surrounding the treatment field and allows local cell-cell and cell-environment interactions to be represented while maintaining computational tractability. Previous experimental work demonstrated that irradiated mammary tissue from immunodeficient hosts exhibits increased macrophage recruitment, persistent inflammation, and enhanced recurrence permissiveness^5,9^. Because recurrence emerges from interactions between immune cells and the evolving tissue microenvironment rather than isolated cellular behaviors, the tissue itself was modeled as a dynamic component of the system^14^. Due to the known role of macrophages at the site^6^, the model was developed as a testing space for elucidating the role of transient immune cell perturbation and downstream persistent inflammation. The model was intentionally designed as the minimal mechanistic representation^19^ capable of testing the hypothesis that persistent inflammatory tissue memory links transient immune remodeling to recurrence. Consequently, fibroblasts, ECM remodeling, and additional cytokines were intentionally omitted to preserve interpretability while testing the proposed mechanism.

### Conceptual Model

Following radiotherapy, mammary tissue undergoes a transient inflammatory response characterized by immune-cell recruitment, macrophage polarization, and cytokine production. Experimental observations show that, in the context of immunodeficiency, inflammation persists well beyond the period expected for normal wound resolution^5,6,8^. Although these observations identify the cellular and molecular components of this recurrent niche, they do not explain how upstream immune perturbation translates into long-term alterations to the tissue, inflammation, and failure to resolve. Therefore, we posit that inflammation becomes integrated as a persistent tissue state that impacts subsequent immune cell behaviors. Conceptually, the model represents recurrence as a feedback process where transient immune behavior modifies the tissue, and the altered tissue subsequently reshapes future immune responses. Model equations explicitly specify the direction of interactions among recurrence associated microenvironment inflammation (RAMI), interleukin-6 (IL6), macrophage phenotype, and tissue permissiveness while population-level macrophage ecology, temporal tissue-state trajectories, and stochastic tumor establishment arise from simulation of the coupled system. The resultant model distinguishes between cellular agents that create tissue remodeling and inflammation and tissue state variables that accumulate and communicate the consequences of these inflammatory events.

### Computational Framework

An ABM was selected because recurrence arises not from the behaviors of one cell or one change in the environment, but from a multitude of coordinated alterations within a specific context. To replicate the complexity of the immune cell and tissue responses, the model framework needs to be capable of representing that heterogeneity. ABMs naturally capture autonomous cell behaviors, local interactions, and emergent tissue-level dynamics. This makes ABMs particularly well-suited for modeling transient immune perturbations and how they result in persistent changes to the tissue state.

The model was implemented in Compucell3D (CC3D), an open-source Cellular Potts modeling platform developed for multiscale tissue simulations. The Cellular Potts model, a lattice-based framework widely used for multi-cellular tissue simulations^21^, hosted in CC3D, represents individual cells as autonomous agents while supporting future incorporation of additional cell populations, diffusible factors, and spatial expansion as the experimental data become available^16^.

### Simulation Execution

Each simulation began with initialization of the irradiated tissue and subsequently advanced through a series of biologically motivated processes performed once per Monte Carlo simulation step (MCS).

#### Initialization

Each simulation began with 50 unpolarized (M0) macrophages randomly distributed among unoccupied lattice sites according to experimentally observed day 0 ratios^6^. Random placement was used to represent macrophages distributed throughout the modeled tissue rather than imposing an initial spatial organization. No tumor cells were present at initialization because the model examines tumor establishment at an irradiated site rather than progression of an existing tumor. RAMI and IL6 were initialized to zero so that inflammatory tissue memory and its downstream mediator emerged from events occurring during the simulation rather than being imposed at baseline.

Simulation time was mapped such that 100 MCS represented one simulated day, and simulations proceeded through day 10 due to experimentally observed tumor cell infiltration^5^. During each simulation, macrophage recruitment, CD4-associated signaling, macrophage phenotype transitions, RAMI accumulation, IL6 emergence, tissue-permissiveness calculation, and tumor-establishment probability were updated sequentially. Model outputs were recorded every 50 MCS and included macrophage abundance and phenotype, RAMI, IL6, tissue permissiveness, and tumor presence.

### Cellular Agents

#### Macrophages

##### Recruitment

Macrophages were initialized as M0 cells and subsequently recruited according to experimentally derived temporal recruitment profiles **(Supplementary Table S1)**^6^. Separate nonrecurrent and recurrent recruitment trajectories were defined using macrophage abundance at days 0, 1, 5, and 10 following radiotherapy. Between experimentally defined time points, target macrophage abundance was estimated by linear interpolation. At each MCS, macrophage abundance was compared with the corresponding target abundance, and additional M0 macrophages were recruited when the simulated population fell below that target. Recruitment per step was limited to prevent abrupt population increases.

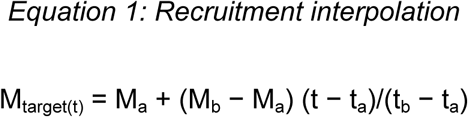

where

- M_target(t)_: target macrophage abundance at time t.
- M_a_: experimentally observed macrophage abundance at the beginning of the interpolation interval.
- M_b_: experimentally observed macrophage abundance at the end of the interpolation interval.
- t_a_: experimental time point corresponding to M_a_.
- t_b_: subsequent experimental time point corresponding to M_b_.
- t: simulated time between t_a_ and t_b_.

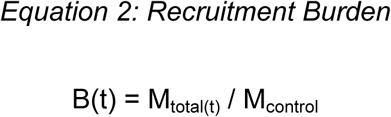

where

- M_total(t)_: current total macrophage abundance.
- M_control_ = 177: day 10 nonrecurrent macrophage reference abundance used to normalize recruitment burden.
- B(t): dimensionless recruitment burden.

##### Macrophage phenotype update

Macrophages occupied one of three phenotypic states: M0 (unpolarized), M1 (anti-tumor), or M2 (pro-tumor). Phenotype switching was stochastic and governed by calibrated transition rates. For a transition from phenotype (P) *i* to P*j*, the per-step transition probability was calculated as

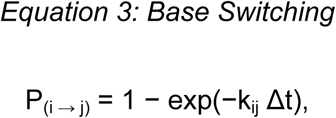

where k__ij_ is the transition rate and Δt=0.1. Baseline transition rates were calibrated prior to introduction of adaptive immune or tissue-mediated feedback.

##### CD4 signaling: Tissue State Variable

CD4+ T-cell effects were represented as global immune-environment presets rather than explicit cellular agents. Previous experimental work from our group identified Th2-skewed CD4+ T-cell crosstalk with macrophage phenotype at the site of recurrence post radiotherapy^17^. These findings motivated representation of CD4-associated immune bias along the Th2 axis in the model. Total CD4-associated signal varied over time while the relative Th2 fraction was determined by the selected immune condition. Because IL4 was the only CD4-associated mediator with active coupling in the reported model, the effective Th2-associated signal was defined as

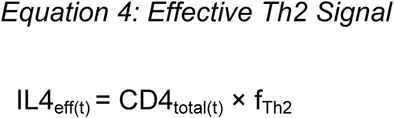

where

- IL4_eff(t)_: effective global Th2-associated IL4 signal at time t.
- CD4_total(t)_: prescribed total CD4 abundance at time t.
- f_Th2_: fixed Th2 fraction for the selected CD4 immune context.

The balanced environment used a Th2 fraction of 0.45, whereas the Th2-biased environment used a fraction of 0.80. The IL4 signal therefore represents a dimensionless effective influence of CD4 composition rather than an experimentally measured cytokine concentration. Although the computational scaffold contains inactive interferon-gamma (IFNγ)- and interleukin-10 (IL10)-like signals, their coupling coefficients were zero and they were not considered active mechanisms in the present study.

In addition, IL4 increased the rates of M0→M2 and M1→M2 polarization. In the final model, IL6 provided an additional M2-directed feedback:

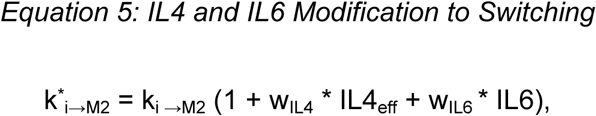

where

- k_i◊j_: calibrated baseline transition rate.
- w*_IL4_* = IL4 coupling coefficient in the final model.
- w_IL6_: IL6-to-M2 coupling coefficient selected by parameter sweep.
- IL6: current emergent tissue IL6 signal.

for *i= M0 o*r *M1*. The IL4 coupling coefficient was fixed at 0.35 following calibration, whereas the IL6 coupling coefficient was selected through the feedback parameter sweep described below. The number of macrophages added during an individual update was limited to the smaller of the population deficit or 0.2M_t_ + 10, where M_t_ represents current macrophage abundance.

##### Recurrence-Associated Microenvironmental Inflammation (RAMI)

RAMI was introduced as a latent state variable representing accumulated unresolved inflammatory remodeling. RAMI integrates three features of the simulated tissue: macrophage abundance relative to the calibrated control, M2-associated remodeling, and the fraction of inflammatory remodeling retained over time. First, macrophage recruitment burden was normalized to the day 10 nonrecurrent control according to **Equation 2**, where M_control_ = 177. M2-associated remodeling incorporated both maintenance by existing M2 macrophages and stronger contributions from newly generated M2 cells:

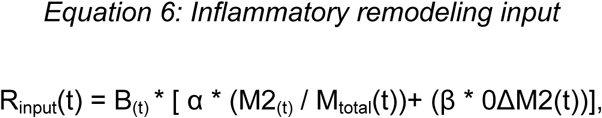

where

- R_input_(t): inflammatory remodeling input at time t.
- α = 0.20: maintenance-remodeling weight.
- β = 1.00: new-event remodeling weight.
- M2_(t)_/M_total_: fraction of macrophages maintaining M2-associated remodeling
- ΔM2: number of newly accumulated M2 macrophages at time t, max[0, M2(t) − M2(t−1)] This distinction allows established M2 macrophages to maintain inflammatory remodeling while newly generated M2 cells represent stronger new inflammatory events. The remodeling contribution was then scaled by the persistence coefficient, which represents the fraction of inflammatory remodeling retained by the tissue.

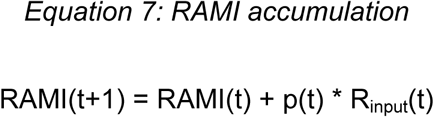

where

- RAMI(t): accumulated recurrence-associated microenvironmental inflammation.
- p(t): time-dependent persistence coefficient determined by the selected persistence condition.
- R_input_(t): inflammatory remodeling input at time t.

Normal-resolution simulations used persistence values of 1.0, 0.8, 0.5, and 0.2 across the modeled temporal intervals, whereas persistent simulations used 1.0, 0.95, 0.90, and 0.85. Thus, both conditions initially retain radiation-associated remodeling but diverge as the normal tissue progressively resolves inflammatory history.

##### IL6 feedback

IL6 was represented as an emergent inflammatory mediator downstream of accumulated tissue remodeling rather than as an independently prescribed input. At each update,

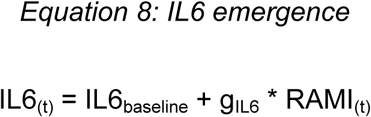

where

- IL6_baseline_ = 0 in the current implementation.
- g_IL6_ = 0.05, gain translating RAMI into the emergent IL6 signal.

The resulting IL6 signal contributed to M2-directed macrophage transition rates as described above, completing the tissue-to-cell component of the feedback loop. In this architecture, RAMI represents accumulated inflammatory history, whereas IL6 represents one experimentally grounded mediator through which that altered tissue state influences subsequent cellular behavior.

### Mechanistic feedback ablation

To distinguish model-encoded relationships from emergent population-level behavior, individual components of the proposed tissue-mediated feedback architecture were selectively ablated while all other model rules and nominal parameter values were held constant. Three model architectures were evaluated under recurrent macrophage recruitment and persistent inflammatory resolution: (1) the complete model, in which RAMI generated IL6 and IL6 modified M2-directed macrophage transitions; (2) a no-feedback model, in which RAMI-derived IL6 remained present but its coupling to macrophage phenotype was removed; and (3) a no-IL6-emergence model, in which RAMI accumulated but did not generate IL6. Each architecture was evaluated across 30 independent stochastic simulations. The day 10 macrophage M2/M1 ratio was designated as the primary outcome for determining whether tissue-to-cell feedback was required to reproduce the experimentally observed recurrent macrophage ecology. RAMI, IL6, tissue permissiveness, and tumor establishment were evaluated as secondary outcomes to characterize downstream consequences of each architectural perturbation.

### RAMI Sensitivity Analysis

To evaluate robustness to hypothesis-driven parameterization, the maintenance-remodeling coefficient (α), new-event-remodeling coefficient (β), persistence scale, RAMI-to-IL6 gain, and IL6-to-macrophage coupling were varied independently across three predefined levels. The resulting 243 parameterizations were evaluated across five stochastic realizations. Robustness of macrophage phenotype was quantified as the absolute distance between simulated day 10 M2/M1 ratios and the experimentally observed recurrent target of 2. These results were used to perform a narrower confirmation experiment. For the confirmation experiment, “high” and “low” refer to the predefined sensitivity bounds, not to values selected after seeing the results, preserving the original interpretation of the screen and avoiding post hoc retuning. To improve interpretability, a single “high” and “low” set were not utilized. Instead, we selected a small set of deliberately different parameterizations that met the following requirements: 1) they were already tested, 2) they were near the recurrent M2/M1 target in the 5-seed screen, 3) they collectively span the low and high bounds of the uncertain parameters, and 4) they remained numerically distinct from each other. This is intended to confirm equifinality under increased stochastic replates, not to further optimize the model.

### Tumor permissiveness

The model represents tumor infiltration and establishment rather than subsequent tumor growth. No tumor cell infiltration was permitted before day 5, allowing the inflammatory microenvironment to develop before evaluation of recurrence permissiveness. Beginning on day 5, the probability of tumor establishment during each update was calculated as

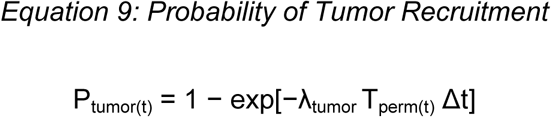

where

- P_tumor(t)_: probability of tumor-cell infiltration during an eligible update.
- λ_tumor_ = 0.001, baseline tumor infiltration rate coefficient.
- Δt = 0.1, model time increment used in the probability conversion.
- T_perm(t)_: tissue permissiveness at time t = w_R_ * RAMI_(t)_ [1 + w_I_ * IL6_(t)_]
- w_R_ = 1.0, RAMI contribution weight.
- w_I_ = 1.0, IL6 amplification weight.

Tumor permissiveness was modeled as a downstream property of both accumulated tissue remodeling and ongoing inflammatory signaling. This formulation gives RAMI a direct contribution to tissue permissiveness while allowing IL6 to amplify the effect of accumulated inflammatory history. Consequently, IL6 alone cannot generate permissiveness in the absence of tissue remodeling. A Bernoulli draw was performed at each eligible timestep. Following successful infiltratoin, a single tumor cell was placed at a randomly selected unoccupied lattice position and no further tumor cells infiltrated. Tumor presence was therefore treated as a binary endpoint reflecting whether the simulated tissue permitted tumor establishment by day 10.

### Parameterization

Model parameters were derived from three sources: direct experimental measurements, calibration to experimentally observed behaviors, and hypothesis-testing parameters introduced to evaluate mechanisms not directly measurable experimentally **(Supplementary Table 1)**. Experimental measurements were prioritized where available while unconstrained parameters were selected through calibration or systematic parameter sweeps rather than assigned to reproduce individual simulation outcomes.

#### Experimentally constrained parameters

Macrophage abundance and phenotype were used as the primary experimental constraints for model development. Nonrecurrent and recurrent macrophage recruitment trajectories were derived from experimentally measured macrophage abundance at days 0, 1, 5, and 10 following radiotherapy^5^, with linear interpolation used to determine target abundance between measured time points. Day 10 macrophage phenotype distributions provided experimental targets for the relative abundance of M0-, M1-, and M2-like macrophages. These measurements established the cellular-scale behaviors against which the baseline model was evaluated prior to introduction of tissue-memory or inflammatory-feedback mechanisms.

#### Calibrated parameters

Baseline macrophage phenotype-transition rates were calibrated to reproduce the experimentally observed macrophage composition under nonrecurrent conditions. Calibration was performed before introduction of CD4-associated IL4 coupling, RAMI-mediated tissue memory, or IL6-mediated feedback so that subsequent mechanisms represented perturbations of an established baseline rather than being incorporated into the initial fit. The calibrated transition rates governed stochastic M0→M1, M0→M2, M1→M2, and M2→M1 transitions and were subsequently held constant during hypothesis-testing experiments. Transition rates were iteratively adjusted until simulated macrophage phenotype distributions reproduced the experimental calibration targets at the evaluated time points.

#### Hypothesis-testing/selected parameters

Parameters without direct experimental equivalents were introduced to test specific mechanistic hypotheses. These included the RAMI maintenance and new-event remodeling weights, normal and persistent tissue-resolution profiles, IL4 coupling strength, and IL6 feedback strength. Rather than interpreting these values as measured biological quantities, each parameter represents the relative strength of a modeled process. Candidate values or formulations were evaluated through targeted simulation experiments, and final values were selected according to predefined behavioral criteria while preserving the experimentally calibrated baseline where appropriate.

### Simulation Experiments

The model was developed incrementally, with each simulation experiment testing whether an additional mechanism was required to reproduce experimentally observed features of the recurrent niche. Unless otherwise stated, all experiments were simulated through day 10 and repeated across 30 independent stochastic realizations.

#### Baseline calibration

The baseline model was first evaluated in the absence of tissue-memory and inflammatory-feedback mechanisms. Macrophage recruitment was constrained by the experimentally derived nonrecurrent recruitment trajectory, and stochastic phenotype-transition rates were calibrated against experimental macrophage phenotype distributions. This experiment established the reference macrophage ecology used for subsequent model perturbations.

#### IL4 coupling

Following baseline calibration, CD4-associated immune bias was introduced through the effective IL4 signal described above. IL4 coupling strength was systematically varied while the calibrated macrophage-transition parameters were held constant. Simulations were performed under balanced and Th2-biased CD4 environments to determine the extent to which adaptive immune bias could alter macrophage phenotype without disrupting the calibrated control state^17^. The selected IL4 coupling coefficient was subsequently fixed for all downstream experiments.

#### Tissue memory formulation hypotheses

Three candidate formulations of inflammatory tissue memory were evaluated to determine how previous inflammatory remodeling should influence future tissue state. Perfect memory retained accumulated inflammatory remodeling without resolution, whereas new-event-only memory represented the tissue state using only newly generated inflammatory events. Mixed memory incorporated both newly generated and previously accumulated remodeling while allowing retention to vary according to the persistence profile. Comparison of these formulations was used to select the tissue-memory architecture for subsequent experiments.

#### Recruitment × persistence

To determine whether experimentally observed recurrence-associated mechanisms were sufficient to reproduce the recurrent tissue state, macrophage recruitment and inflammatory persistence were varied independently in a 2 × 2 factorial design. Simulations combined either nonrecurrent or recurrent macrophage recruitment with either normal or persistent inflammatory resolution, producing four conditions: nonrecurrent/normal, nonrecurrent/persistent, recurrent/normal, and recurrent/persistent. The experiment was performed under both balanced and Th2-biased CD4 environments. Macrophage M2/M1 ratio and RAMI were evaluated as cellular- and tissue-level outcomes, respectively.

#### IL6 feedback

Finally, tissue-to-cell inflammatory feedback was introduced by coupling RAMI-derived IL6 to M2-directed macrophage phenotype transitions. The IL6 coupling coefficient was systematically varied while all previously selected model parameters were held constant. Candidate coupling strengths were evaluated under both calibration and recurrence-associated conditions to determine whether inflammatory feedback could reinforce macrophage phenotype without substantially disrupting the calibrated nonrecurrent ecology.

### Statistical Analysis

Unless otherwise stated, simulation results are reported as mean ± standard deviation across 30 independent stochastic realizations. Sensitivity analyses evaluated robustness across biologically plausible parameter ranges. Inferential statistics were reserved for testing mechanistic hypotheses, whereas descriptive statistics and regression analyses were used to characterize emergent model behavior.

### Modeling Assumptions

To represent the tissue while also preserving interpretability, certain reasonable assumptions were made. Here, the tissue is represented in a 2D cross-section where cells occupy equal lattice volume. Our model does not study the progression of recurrence, but its incidence, so we use a binary to represent whether tumor cells arrive at the site. Inflammation as a product of transient immune alterations required development of a receiver, repository, and transmitter of inflammatory events. RAMI is a latent state variable representing the accrual of inflammation over the course of wound-healing as opposed to a quantifiable biological factor. RAMI is upstream of the mediator and consequence of persistent inflammation IL6. IL6 has been found in our work to increase M2/M1 ratio and increase tumor cell invasion^6^, so it is an experimentally grounded choice. Fibroblasts and ECM were omitted here to maintain the simplest representation of the tissue to improve interpretability.

## Results

### The base model replicates experimentally observed macrophage infiltration and the recurrence-resistant phenotype

Our site of interest is a highly heterogeneous microenvironment that becomes increasingly complex during acute damage to normal mammary tissue^2^. ABMs are uniquely poised to represent such a site^18,19^. This is in part due to the autonomy of the agents, which allows them and their environment to develop emergent behaviors^16^. We aim to use our model to investigate the intersection of experimentally observed cellular relationships, environmental factors, and tissue characteristics in a controlled, interpretable framework.

Here, we represent the irradiated tissue section as a lattice seeded with macrophages **(Supplementary Table 1)**. Prior to hypothesis testing, we first ensure that the model can be calibrated to replicate the nonrecurrent environment. Previous work has found that in nonrecurrent mice, macrophage composition is close to 90% M0, 5% M1, and 5% M2^6^. First, we constrained macrophage recruitment to match experimental immunohistochemistry data (**Figure 3A**^5^**; Equation 1, 2)**. Once it was confirmed that macrophage recruitment met experimentally proportional population sizes at days 0, 1, 5, and 10 days post-radiation, we then employed base polarization and phenotype switching equations for macrophages **(Equation 3)**. Using experimental data of macrophage composition at day 10 post-radiotherapy, we iteratively adjusted the rate constants until we reliably met the expected range: 88.5±1.76% M0, 5.33±1.77% M1, and 6.17±1.44% M2 (**Figure 3B, C)**. The model therefore replicated experimental observations of macrophage recruitment and population composition, providing a calibrated baseline for subsequent hypothesis testing.

**Figure 3.**
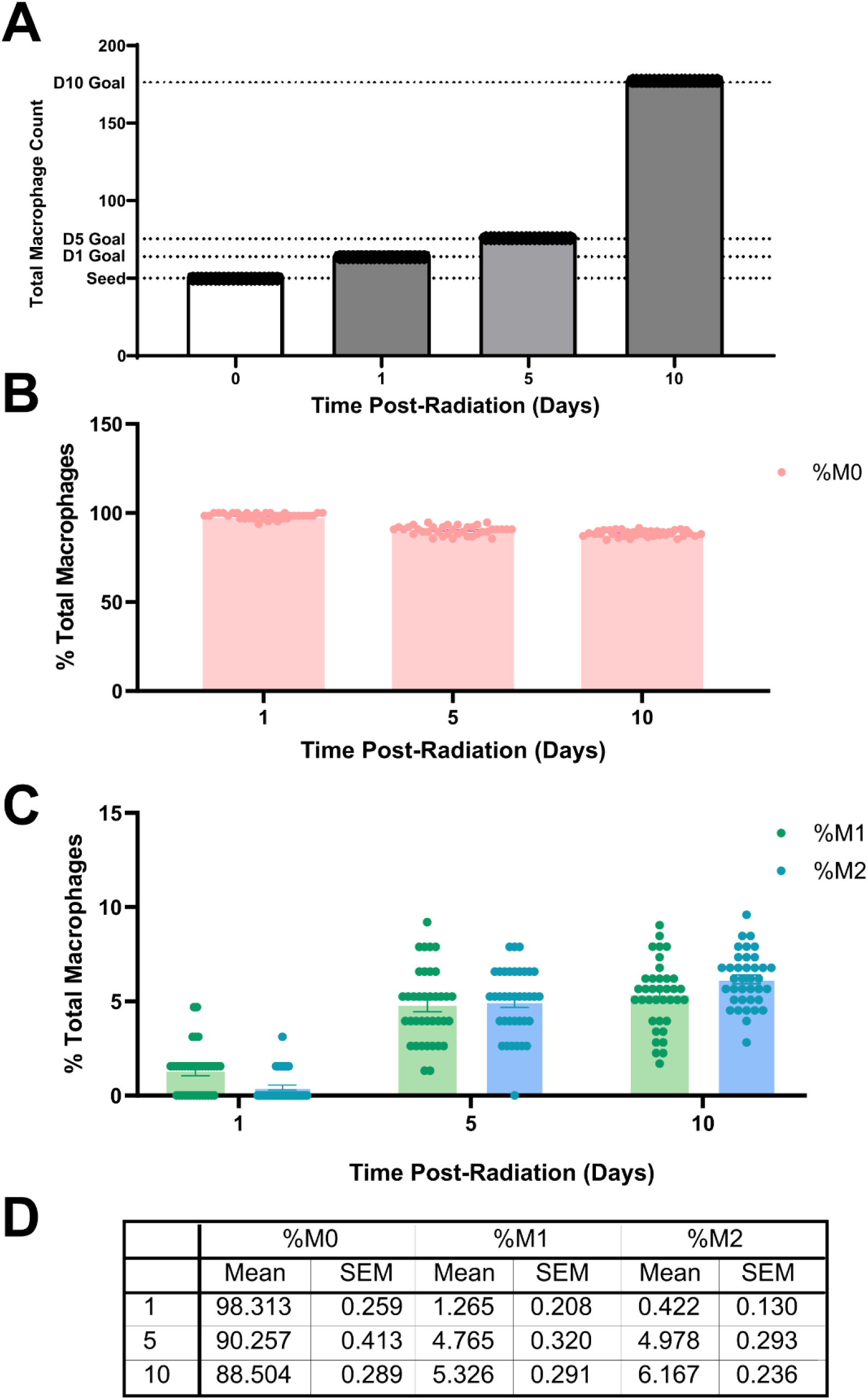
The calibrated model reproduces experimentally observed macrophage characteristics. (A) Simulated macrophage recruitment matches experimentally observed macrophage numbers at 0, 1, 5, and 10 days post-radiotherapy. Simulated macrophage phenotypes M0 (B), M1 (C), and M2 (C) meet expected % macrophage compositions. (D) Mean ± SD macrophage composition at days 1, 5, and 10 post-radiotherapy across 30 stochastic simulations.

### CD4+ T-cells alone cannot reshape macrophage ecology in the context of recurrence-resistance except at extreme cytokine coupling

Due to the well-established relationship between macrophages and CD4+ T-cells^22,23^, we sought to establish the sensitivity of the system to CD4+ T-cell-derived IL4, a Th2 cytokine that causes macrophages to adopt an M2-like phenotype. Macrophage plasticity allows them to respond to the environment, which we bias using CD4+ T-cell presets as shown in **Equation 4**.

Assuming that the calibrated system implicitly captures the overall impact of IL4, we determined how sensitive the system is to changes in CD4 T-cell-derived IL4 at the site (**Figure 4A**). We applied **Equation 5** and systematically varied IL4 coupling weights to find the largest, biologically reasonable coupling coefficient that preserves the experimentally calibrated control under balanced CD4 conditions but increases M2/M1 in the Th2 bias condition. This allowed us to observe the impact of CD4-T-cell-derived IL4 in the system while not allowing it to control or override the calibrated behaviors in the system. The effect of IL4 on macrophage phenotype was moderate. M1 and M2 percentages did diverge, but only at high coupling coefficients sufficiently large to override the calibrated baseline did the balanced (**Figure 4B**) or Th2-biased (**Figure 4C**) environment capture the M2/M1 ratio of approximately 2.

**Figure 4.**
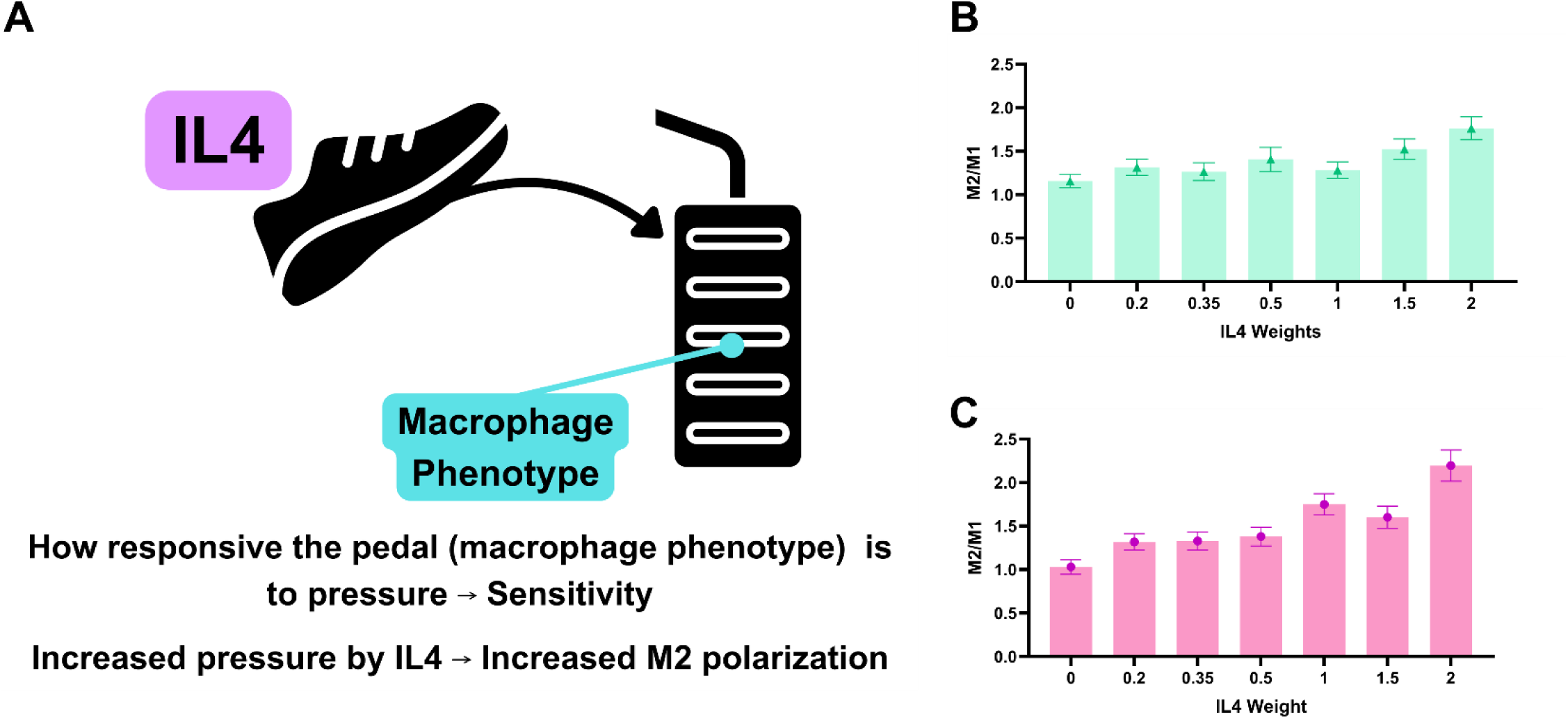
Sensitivity of macrophage polarization to CD4-derived IL4. (A) Schematic illustrating the IL4 coupling coefficient. (B) M2/M1 ratio as IL4 coupling increases under balanced CD4 conditions. (C) M2/M1 ratio under Th2-biased CD4 conditions. Error bars represent SD from 30 stochastic simulations.

To choose a coupling weight that best represents the non-recurrent environment and does not risk overtaking other relationships in the model, we chose the highest IL4 coupling coefficient that did not perturb the control as shown in **Figure 4C**. The responsiveness or sensitivity of macrophage ratio to IL4 can be broken into 3 regions: low (0-0.35), transition (0.5-1), and high (>1). To preserve the control, we chose the IL4 coupling coefficient 0.35, which resulted in a day 10 post-radiotherapy M2/M1 of 1.27 ± 0.387. In the Th2 biased condition, we see a slightly noisier response, which is to be expected as Th2 bias allows for increased strength of IL4 signaling. CD4+ T-cells could shift the macrophage balance toward recurrent macrophage ecology, but only at the highest IL4 coupling coefficients, even in Th2-biased environmental models. This implies that there is a layer of biology that is not represented in the model, decreasing the probability that the macrophage ecology of the site can be perturbed to reflect recurrent site characteristics.

### RAMI is a state variable that represents how inflammation accumulates

Persistent inflammatory signaling has been observed experimentally following radiotherapy, suggesting that recurrence may reflect a failure of tissue resolution rather than a prolonged acute response. To that end, we have established the presence of inflammatory cytokines present beyond their usual timeline in the recurrent wound^6^. This and other work suggest that the recurrent site is not simply wounded, it is persistently inflamed due to aberrant resolution of inflammation^5,6,9^. We hypothesize that this inability to resolve inflammation is the force that would cause the site to undergo a state shift from nonrecurrent to recurrent, as represented by the macrophage ecology. For there to be unresolved inflammation, there must be memory of inflammation.

The control tissue machinery calibrated previously will be subject to 3 hypotheses about tissue memory at the site to determine what best represents how the tissue remembers inflammatory events: 1) In perfect memory, this tissue is inflicted and never forgets inflammatory events, accruing them similarly to how we hypothesize tissue inflammation persists at the site; 2) in new event memory, the site is impacted only by new inflammatory events at the site, assuming resolution does occur as time goes on; and 3) in mixed memory, the tissue maintains memory of inflammatory events according to a coefficient that determines how capable the tissue is of resolving inflammation. We present RAMI as a mechanistic abstraction to represent the latent tissue state and reflect accrued inflammatory information. RAMI is both receiver and transmitter, storing inflammatory event information and then communicating it downstream. The 3 competing hypotheses were evaluated to determine how irradiated tissue retains inflammatory information to decide what RAMI stores.

Injured tissues resolve inflammation. Therefore, a model that never forgets inflammatory events cannot represent normal tissue repair. Perfect memory (**Figure 5A**) is biologically unrealistic as the tissue will go through some amount of resolution. The difference between the nonrecurrent and recurrent sites is that one resolves inflammation and the other does not. Here, RAMI, the latent state variable representing unresolved, accumulated inflammation, rises into extreme numbers by day 10 post-radiotherapy. This models an environment where resolution does not occur, which removes one of the fundamental behaviors of injured tissue. Perfect memory does provide us with an illustration of the upper, most extreme bounds of RAMI. For this reason, perfect memory captures an upper bound on inflammatory accumulation but does not represent physiologic resolution. In contrast, new event memory does (**Figure 5B**) the opposite of the perfect memory model. While its retention of inflammation is decreased, it is now miniscule. While the concept is more biologically relevant, it now lacks accrual. There is no accrual of inflammation from previous time points and no mechanism for resolution due to lack of memory. Finally, mixed memory (**Figure 5C; Equation 6, Equation** 7) accounts for both perfect memory and new event only memory, resulting in moderate RAMI values (11.58 ± 3.26). Because RAMI is a latent model state rather than a directly measurable biological quantity, candidate memory formulations were evaluated according to their qualitative behavior: whether they permitted accumulation, resolution, and divergence between normal and persistent inflammatory conditions rather than against an absolute experimental RAMI value. The system takes new events into account to a greater extent than events from previous timepoints, but this is scaled by a coefficient called persistence, which determines how impaired resolution of the tissue is. At high values, persistence minimizes the effects of resolution of inflammation. At low values, persistence allows resolution to dampen new inflammation, reducing what is stored from one time point to the next. This is the version of tissue memory we selected for advancement. The tissue’s ability to remember inflammation can be captured by a latent state variable that can receive inflammatory signal and transmit those aggregate impacts downstream. This tissue-memory representation becomes the foundation for subsequent experiments investigating the transition from inflammatory persistence to recurrence.

**Figure 5.**
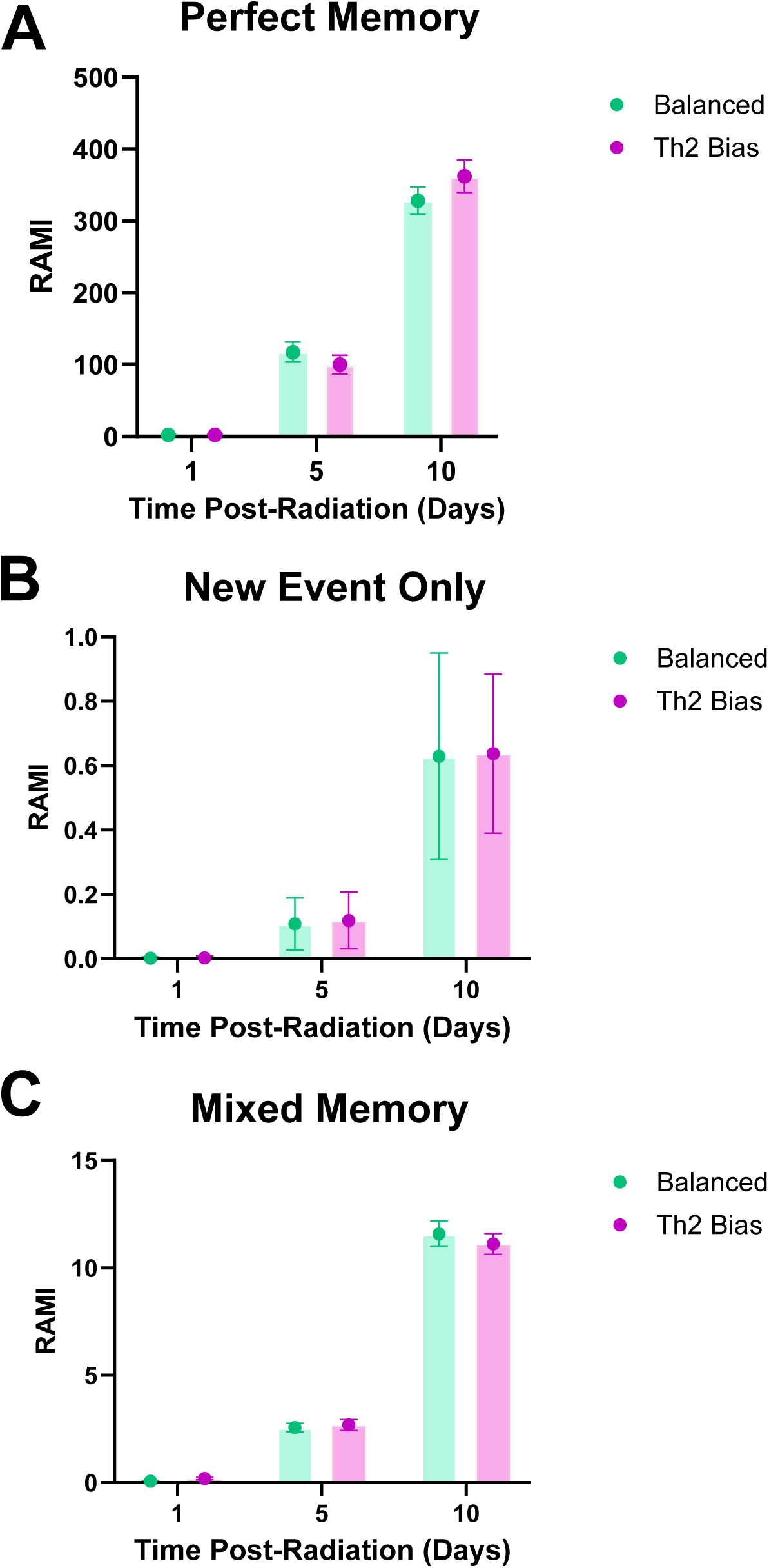
Evaluation of candidate tissue memory formulations. RAMI accumulation over 10 days following radiotherapy using perfect memory (A), new event only memory (B), and mixed memory (C) formulations in balanced and Th2-biased environments. Error bars represent SD from 30 stochastic simulations.

### Increased macrophage recruitment and inflammatory persistence are insufficient to replicate the recurrent macrophage ecology but do replicate accumulating inflammation

It is at this stage that it is important to understand the impact of the changing environment on RAMI. We dissected recurrence into 2 experimentally observed mechanisms: 1) persistent inflammation, and 2) high macrophage recruitment to determine whether one or both could explain the recurrent microenvironment. The nonrecurrent context will be challenged by: 1) applying the experimentally observed recurrent macrophage recruitment curve, and/or 2) increasing the tissue’s inflammatory memory via persistence. This will allow us to determine if either the experimentally observed change in macrophage abundance or the tissue’s inability to resolve inflammation are independently responsible for the shift from the nonrecurrent to the recurrent state. This would be confirmed by increased RAMI and an increase in M2 macrophages over M1 macrophages at a ratio of about 2:1.

Though the maximum increase of the M2/M1 ratio was approximately 1.75 (**Figure 6A, B)** in both balanced and Th2-bias regardless of recurrent recruitment or persistence (**Figure 6C, D)**, we see that recurrent recruitment and persistence together increase RAMI values (**Figure 6E, F)** to approximately double the normal tissue controls. Recruitment significantly altered RAMI (F(1,116) = 204.3, p<0.0001), as did persistence (F(1,116) = 171.2, p<0.0001), with a significant recruitment × persistence interaction (F(1,116) = 17.3, p<0.0001). Recruitment and impaired resolution were the dominant contributors to variation in RAMI in both immune contexts (35–40% and 33% of the total variation in the ANOVA decomposition, respectively). Although their interaction is statistically significant, it accounts for a comparatively small proportion of the variation (3–5%), indicating that both mechanisms independently contribute to inflammatory tissue memory.

**Figure 6.**
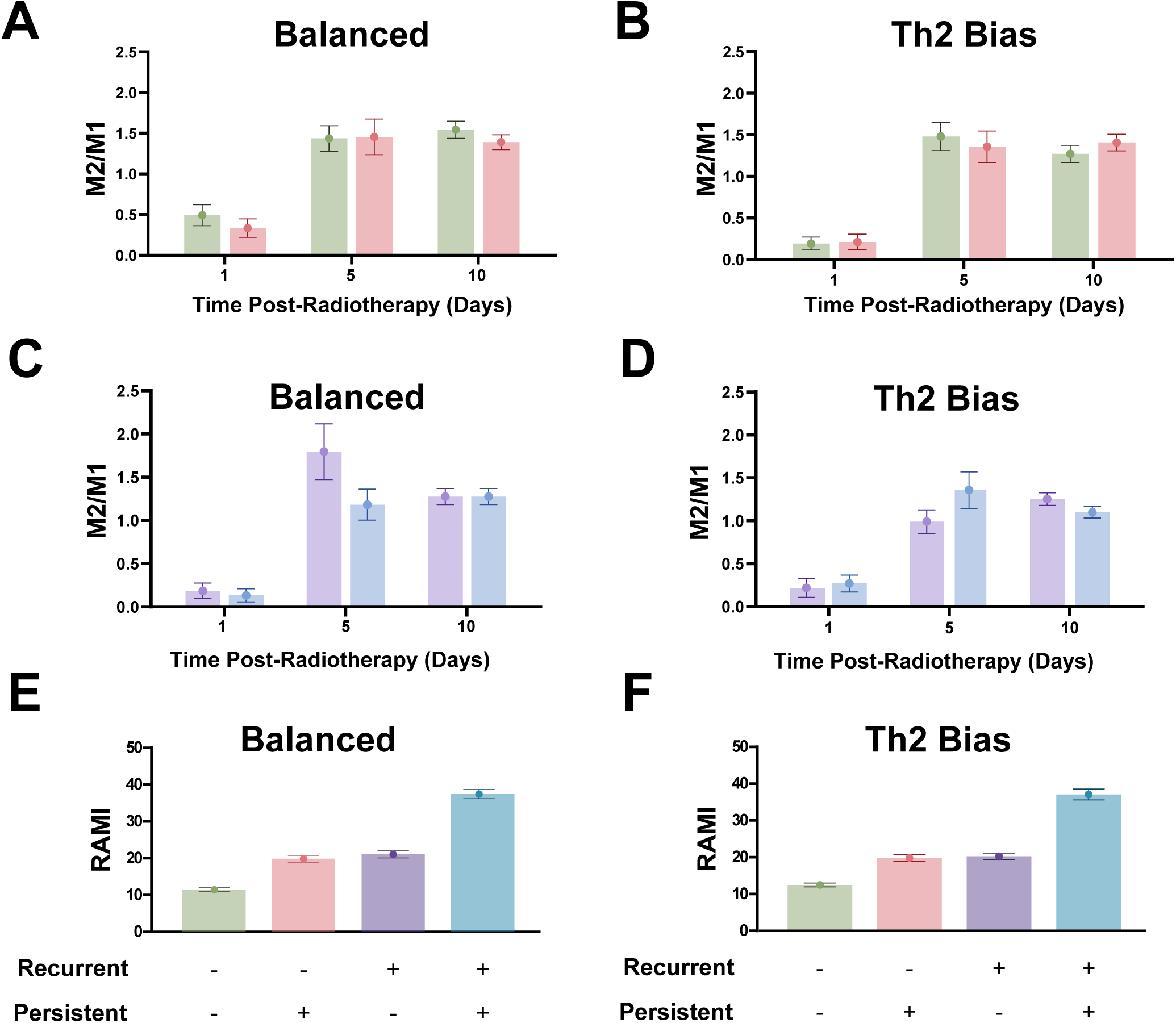
Recruitment and impaired inflammatory resolution jointly increase inflammatory tissue memory. M2/M1 ratios under normal and persistent inflammatory resolution are shown for nonrecurrent macrophage recruitment in balanced (A) and Th2-biased (B) CD4+ T cell environments and for recurrent macrophage recruitment in balanced (C) and Th2-biased (D) CD4+ T cell environments. Day 10 RAMI under combinations of nonrecurrent or recurrent macrophage recruitment and normal or persistent inflammatory resolution is shown in (E) balanced and (F) Th2-biased CD4 environments. Data represent mean ± SD from 30 independent stochastic simulations.

Recruitment accounts for the largest proportion of the variation in the M2/M1 ratio under balanced conditions (43%), whereas persistence contributes a smaller but significant effect (7%). In Th2-biased environments, recruitment and persistence each exert substantial independent effects (35% and 27%, respectively), while their interaction is not significant. Persistence fails to reproduce the recurrent macrophage ecology. The same is true of increased macrophage recruitment as it is insufficient to produce a model of the recurrent macrophage composition. When the site experiences both the recurrent macrophage recruitment curve and high persistence, RAMI more than doubles, indicating that the site is accruing inflammation as expected, so the model requires other biological factors that would allow it to reproduce the recurrent environment.

### IL6 feedback provides the missing layer of biology that recapitulates recurrent macrophage ecology

Inflammatory tissue memory may require an explicit feedback mechanism to reproduce the recurrent macrophage ecology. Our previous work found that irradiated normal tissue has a profound impact on recruited macrophages. However, we chose not to model the ECM directly. This is because there are a handful of factors identified at the site that have been identified as pivotal for macrophage skewing toward an M2-like phenotype. IL6 is a pleiotropic cytokine typically known for being pro-inflammatory^24–26^. It is associated with increased M2/M1 at the site as well as tumor cell infiltration of tissues^6^. To avoid the assumption that IL6 must come from the macrophages and to appreciate the impact of the tissue, IL6 is generated from the environment as a tissue-level variable **(Equation 8)**. It is cued by RAMI and then impacts macrophages according to the IL6 coupling weights and increases the likelihood of tumor cell infiltration. IL6 coupling coefficients were systematically varied under calibration (nonrecurrent) and validation (recurrent recruitment + persistent inflammatory resolution) conditions. We observe the impact of the macrophage recruitment curve, persistence coefficient, and range of IL6 coupling weights on macrophage composition, M2/M1 ratio, and RAMI to determine: 1) the smallest an IL6 coefficient which preserves the control, nonrecurrent tissue properties while producing biologically meaningful reinforcement under recurrent conditions; 2) if RAMI responds and rises accordingly with no exaggerated read-outs; or 3) if IL6 requires both recurrent macrophage recruitment and persistent inflammatory signaling to maintain the recurrent site characteristics. IL6 can increase M2/M1 to approximately 2, even at moderate coupling weights (where IL6 coupling coefficient=1, M2/M1 = 2.09 ± 0.55) in the context of nonrecurrent tissue (**Figure 7A**). To preserve the control, IL6 coupling coefficient, 0.75 was selected as the largest coefficient that replicated baseline behavior while producing recurrence-associated reinforcement (M2/M1 = 1.62 ± 0.48). When in the recurrent tissue context (**Figure 7B**), coupling weights that preserved the nonrecurrent control met or exceeded the experimentally observed recurrent macrophage ecology (M2/M1 = 3.39 ± 0.73). Moderate IL6 feedback is sufficient to reinforce the recurrent phenotype while preserving the calibrated control. IL6 signaling is generated in the normal tissue (**Figure 7C**) with relatively little change across IL6 coupling weights. However, RAMI-derived IL6 signal is generated with a steady increase across IL6 coupling coefficients in the recurrent tissue (**Figure 7D**). These trends align with RAMI (day 10 post-radiotherapy = 13.53 ± 3.39) (**Figure 7E, F)**, establishing the necessary mechanisms to recapitulate the nature of the tissue.

**Figure 7.**
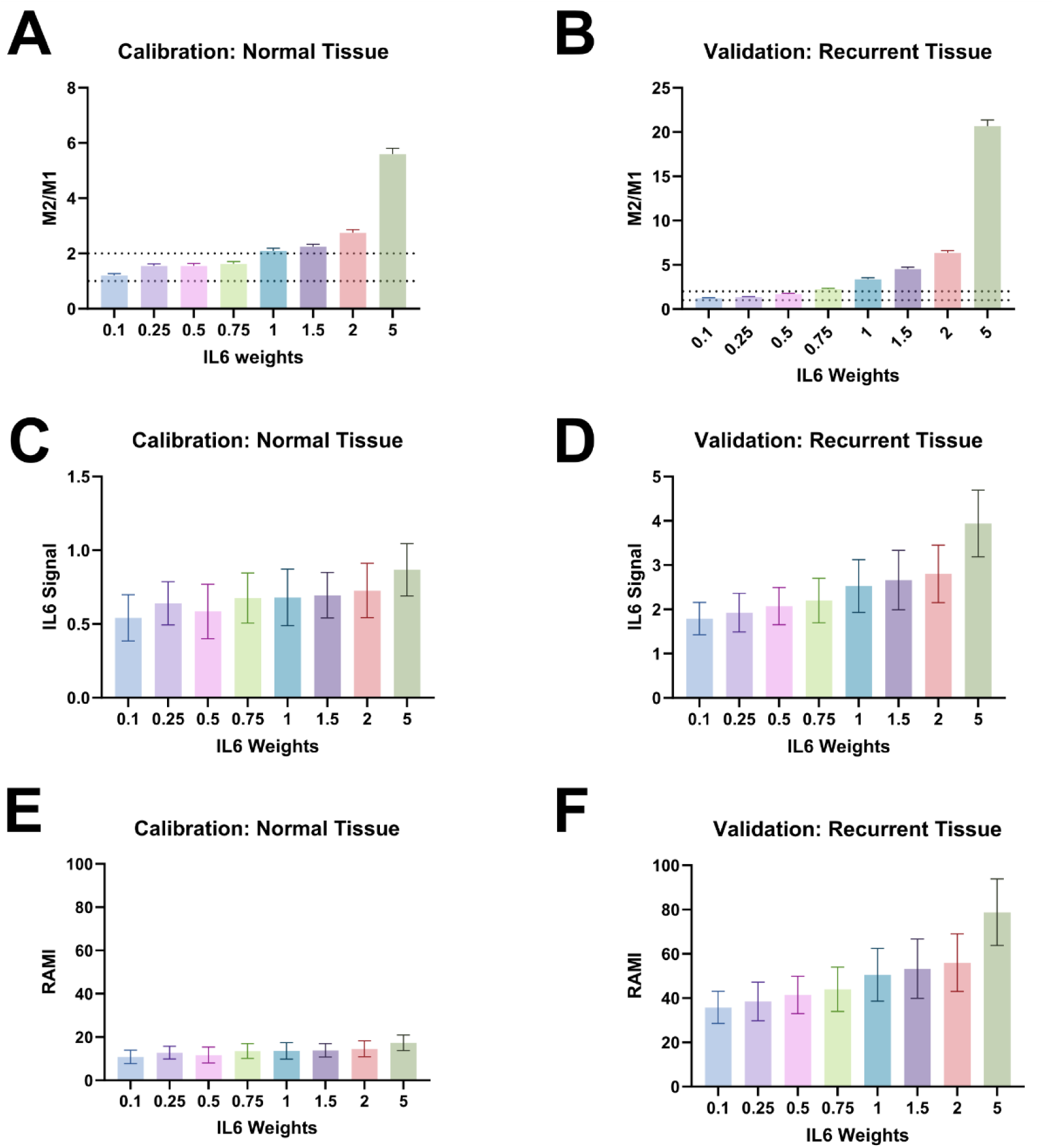
Calibration and validation of IL6-mediated inflammatory feedback. IL6 coupling coefficients were evaluated in calibration (left) and validation (right) environments. (A, B) M2/M1 ratio. (C, D) IL6 signal. (E, F) RAMI. Calibration was performed under nonrecurrent conditions, whereas validation used recurrent recruitment with persistent inflammatory resolution. Bars represent mean ± SD from 30 stochastic simulations.

### Tissue-to-cell inflammatory feedback is required to reproduce recurrent macrophage ecology

We next determined the biological consequences of the tissue being unable to resolve inflammation after radiotherapy. With the IL6 weight now fixed as was previously described, the normal tissue and recurrent tissue ran for 10 days post-radiation. At 10 days (1000 MCS), the tissue was evaluated for macrophage ratio, RAMI, and whether or not tumor cells infiltrated **(Equation 9)**. At day 10 (1000 MCS), the recurrent condition maintained a higher M2/M1 ratio than the nonrecurrent condition (nonrecurrent: 1.63 ± 0.48; recurrent: 2.25 ± 0.55; **Figure 8A**). Recurrent tissue also exhibited greater RAMI (nonrecurrent: 13.53 ± 3.39, recurrent: 44.012 ± 10.02; **Figure 8B**), IL6 signal (nonrecurrent: 0.68 ± 0.17, recurrent: 2.2 ± 0.5; **Figure 8C**), and probability of tumor cell infiltration (nonrecurrent: 33%, recurrent: 60%; **Figure 8D**) than the nonrecurrent tissue. Together, these outputs demonstrate that the completed model differentiates the experimentally defined nonrecurrent and recurrent tissue contexts. However, because IL6 is explicitly coupled to RAMI, IL6 increased with RAMI as expected. We therefore evaluated whether closing this feedback altered the emergent macrophage population state.

**Figure 8.**
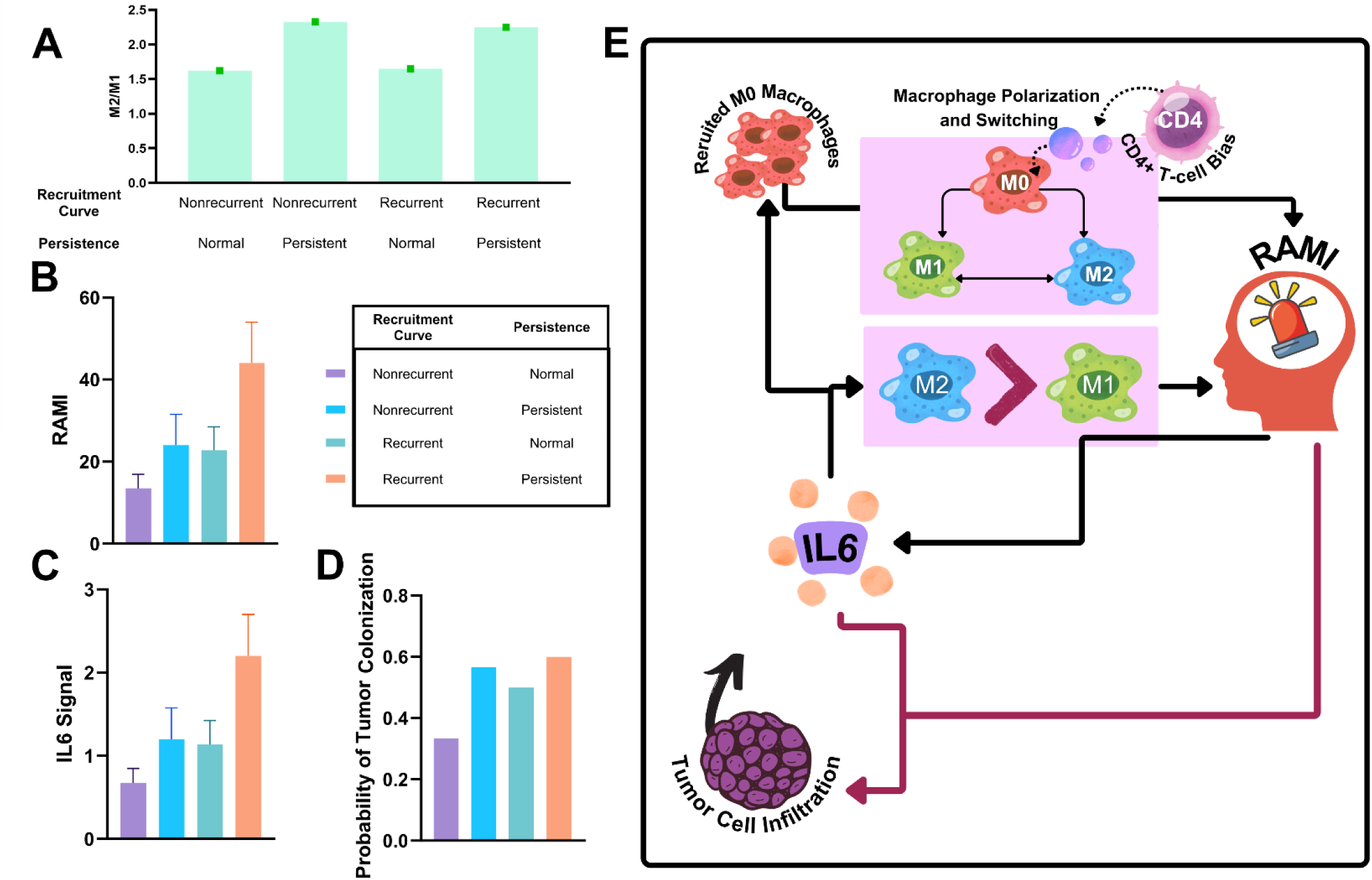
Consequences of inflammatory tissue memory in the recurrent niche model. (A) M2/M1 ratio. (B) RAMI. (C) Emergent IL6 signal. (D) Probability of tumor colonization. (E) Schematic of the tissue-memory feedback architecture linking macrophages, RAMI, IL6, and tumor cell infiltration. Bars represent mean ± SD from 30 stochastic simulations.

### Closing the macrophage-tissue state variable loop is necessary to reproduce the recurrent phenotype

IL6 is explicitly defined as a function of RAMI, while tissue permissiveness and tumor-establishment probability are specified downstream of RAMI and IL6. The population-level macrophage response, in contrast, arises from stochastic phenotype transitions within the coupled system. We therefore next tested whether closing the RAMI-IL6-macrophage feedback loop was required for the recurrent macrophage ecology to emerge. To separate the effects of accumulated inflammatory tissue memory from tissue-to-cell feedback, individual components of the feedback architecture were selectively ablated while all other model parameters were held constant (**Figure 9A**). In the full, no-ablation model, the day 10 M2/M1 ratio reached 2.93 ± 1.58. Preventing RAMI-derived IL6 from impacting macrophage phenotype reduced M2/M1 to 1.16 ± 0.43, despite continued RAMI accumulation and IL6 generation (**Figure 9B**). Similarly, preventing RAMI from generating IL6 reduced M2/M1 to 1.07 ± 0.48. Feedback ablation also reduced RAMI from 43.28 ± 12.62 in the complete model to 35.72 ± 6.95 when IL6-to-macrophage coupling was removed and 35.17 ± 8.79 when IL6 emergence was ablated (**Figure 9C**). Because RAMI accumulation was not directly disabled in either condition, this reduction reflects loss of feedback reinforcement through altered macrophage population behavior. This shows that accumulation of inflammatory tissue memory alone was insufficient to reproduce the recurrent macrophage ecology. Instead, the tissue state had to be communicated back to the macrophage population to close the feedback loop.

**Figure 9.**
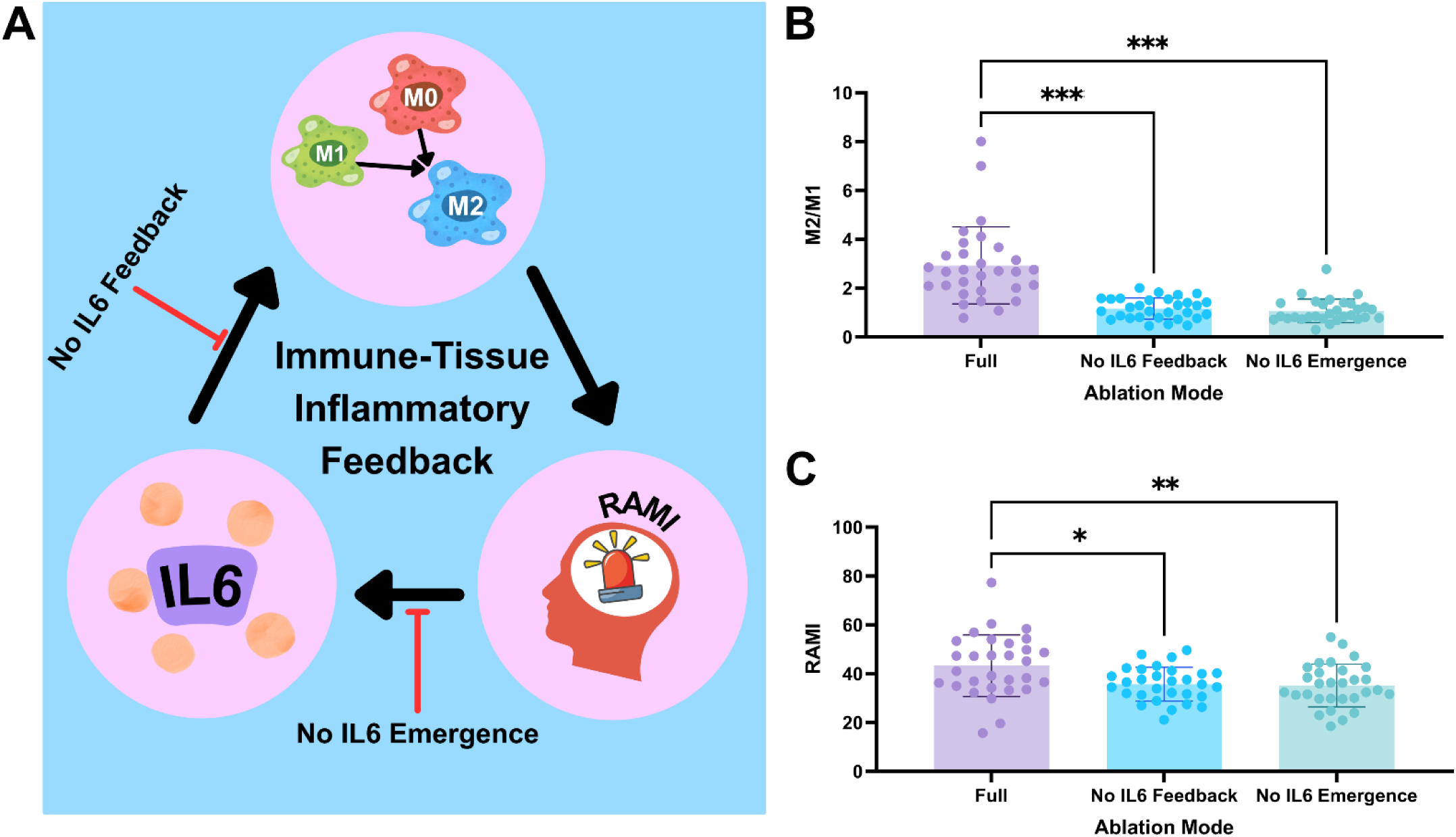
Tissue-to-cell inflammatory feedback is required to reproduce the recurrent macrophage ecology. (A) Individual components of the RAMI–IL6–macrophage feedback architecture were selectively ablated while all other model parameters were held constant under recurrent recruitment and persistent inflammatory conditions. (B) Day 10 M2/M1 ratios for the complete model (Full), a model in which RAMI-derived IL6 does not feed back onto macrophage phenotype (No IL6 Feedback), and a model in which RAMI does not cue IL6 generation (No IL6 Emergence). (C) Day 10 RAMI across the same model architectures. Each point represents an independent stochastic simulation (n=30 per condition). Data are shown as mean ± SD. Statistical significance was evaluated by one-way ANOVA with *p<0.05, **p<0.01, and ***p<0.001.

### Recurrence-associated macrophage behavior can be generated by a family of parameter combinations

To determine whether the recurrent macrophage phenotype depended on the nominal values of hypothesis-driven parameters, we varied the maintenance-remodeling coefficient (α), new-event-remodeling coefficient (β), persistence scale, RAMI-to-IL6 gain, and IL6-to-macrophage coupling across structured ranges around the nominal parameterization. The primary robustness outcome was fixed before analysis as the absolute distance from the experimental recurrent M2/M1 target of 2: D_target_ = |M2/M1 - 2|, where lower values indicate closer agreement with the experimental target.

An exploratory five-seed experiment of 243 parametrizations identified several parameter sets that reproduce recurrent macrophage ecology. Five contrasting near-target parameterizations and the nominal parameterization were subsequently evaluated across 30 independent stochastic realizations. The selected parameterizations spanned the tested ranges of maintenance remodeling, new-event remodeling, inflammatory persistence, IL6 gain, and IL6-to-macrophage coupling (**Figure 10A**). Multiple distinct parameter combinations produced macrophage phenotypes close to the experimentally observed recurrent target. At day 10, the nominal parameterization produced an M2/M1 ratio of 2.63 ± 1.13. Mean M2/M1 ratios for the alternative parameterizations were 1.81 ± 0.90 (A), 1.82 ± 0.70 (B), 1.83 ± 0.70 (C), 1.88 ± 0.79 (D), and 2.24 ± 1.03 (E) (**Figure 10B**). Mean absolute distance from the experimental target ranged from 0.58 to 0.76 among the alternative parameterizations, compared with 0.83 for the nominal parameterization.

**Figure 10.**
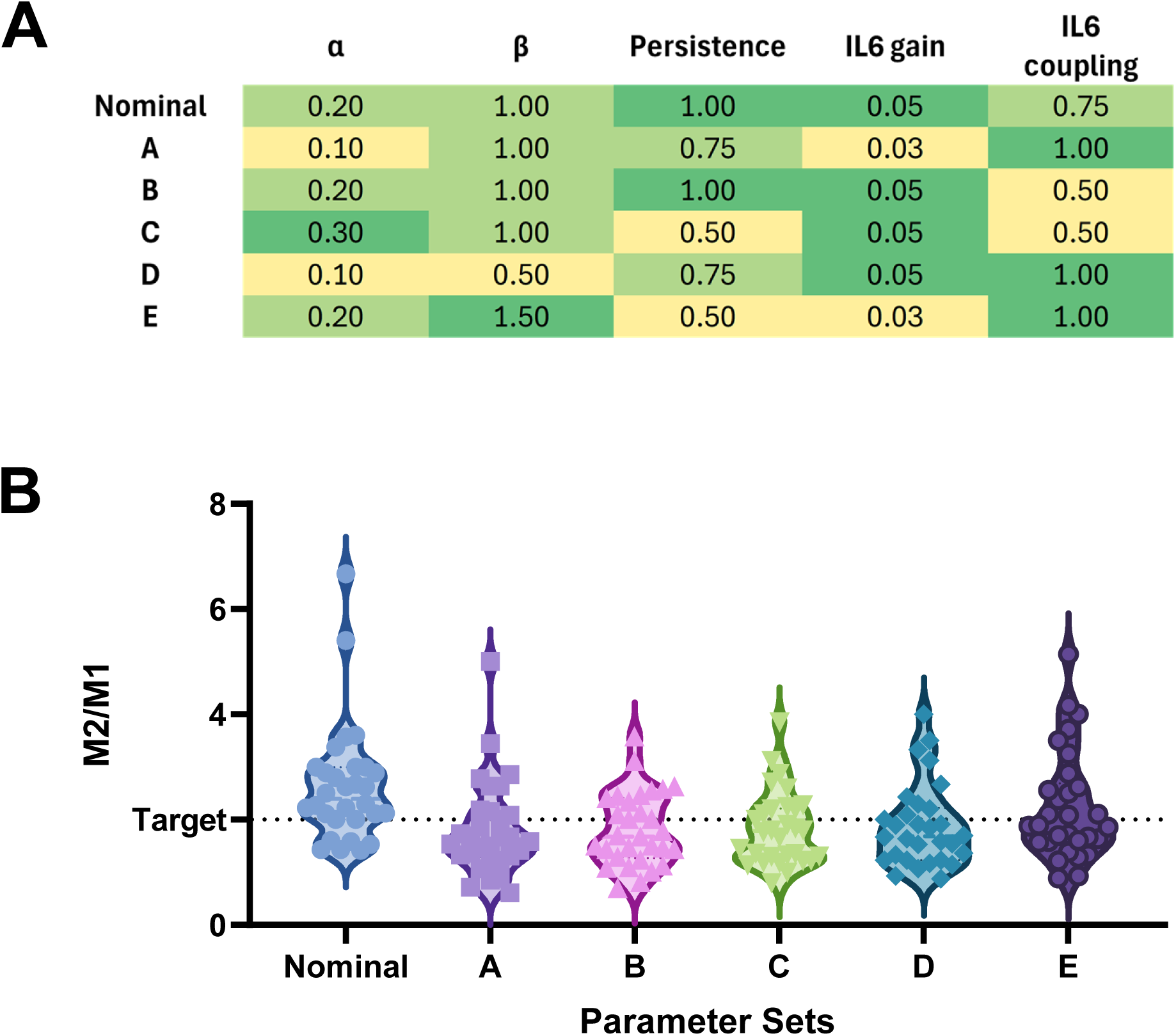
Multiple parameter combinations produced macrophage phenotypes that approximate the experimentally observed recurrent target. (A) Parameter values for the nominal model and five contrasting parameterizations selected from an initial sensitivity screen of 243 combinations. Parameters include the maintenance-remodeling coefficient (α), new-event-remodeling coefficient (β), inflammatory persistence scale, RAMI-to-IL6 gain, and IL6-to-macrophage coupling. (B) Day 10 M2/M1 ratios for each parameterization following higher-replicate confirmation. Each point represents an independent stochastic realization (n=30 per parameterization). Data are shown as mean ± SD. The dashed horizontal line denotes the experimentally observed recurrent M2/M1 target of 2.

## Discussion

We developed an experimentally constrained ABM to investigate how transient immune remodeling after radiotherapy becomes persistent inflammatory tissue memory. Neither increased macrophage recruitment nor impaired inflammatory resolution alone were sufficient to reproduce the recurrent microenvironment. Instead, a recurrence-permissive tissue state emerged through the interaction of inflammatory tissue memory and tissue-mediated feedback, providing a mechanistic explanation for persistent inflammatory signaling and increased tumor permissiveness (**Figure 11**).

**Figure 11.**
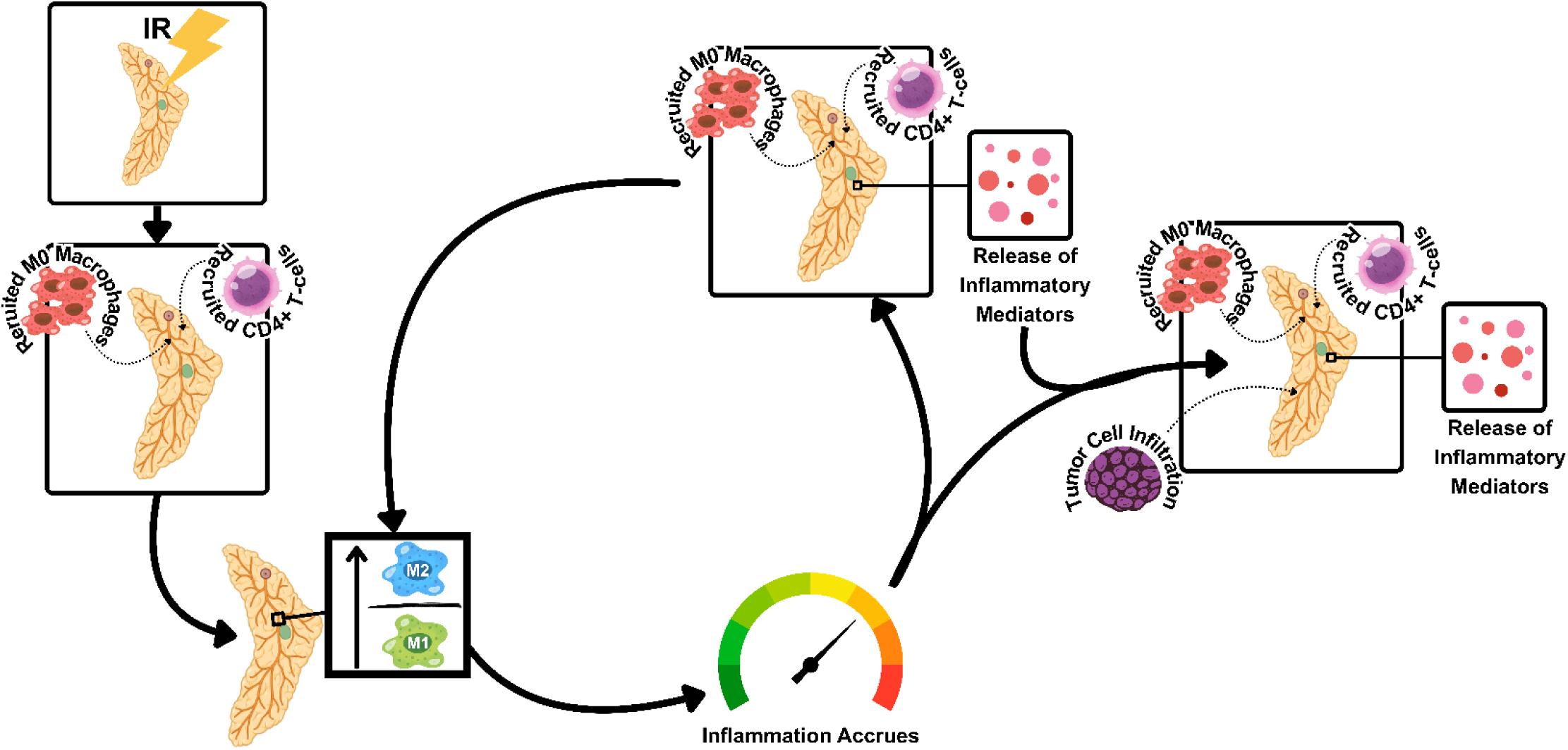
Proposed mechanism linking transient immune remodeling to recurrent tumor permissiveness. Following radiotherapy, transient immune perturbations alter macrophage recruitment and phenotype, resulting in accumulation of inflammatory tissue memory. Persistent tissue-state remodeling promotes inflammatory mediator release, reinforcing macrophage polarization and increasing tumor cell infiltration.

### Failed inflammatory resolution, rather than inflammatory magnitude alone, is important to recurrence

Our results suggest that the magnitude of inflammatory insult alone is insufficient to fully explain recurrent niche development. While increased macrophage recruitment based on experimentally-observed results in the recurrent context did substantially increase RAMI, the tissue’s ability or lack thereof had an independent impact on RAMI. This suggests that the recurrent context requires both increased macrophage recruitment and inability to resolve the wound to synergistically increase accumulation of inflammation. Still, the tissue’s recurrent niche is incomplete in that the experimentally-observed macrophage ecosystem fails to emerge. In retrospect, this is unsurprising as RAMI lacked crosstalk with macrophages. It fails to represent biology sound as tissue-level factors are vital in macrophage plasticity^18,27^. This is consistent with computational and experimental models of wound healing in which the duration and resolution of inflammatory signaling, rather than inflammatory activity alone, determine whether tissue returns to homeostasis or progresses toward chronic pathology^19,28–30^.

Key experimental work found that tumor cell infiltration was preceded by an influx of macrophages^5^, leading to remodeling of the site. Subsequent work identified IL6 as a central inflammatory mediator associated with M2-like macrophage polarization and increased recurrence^6,24,31–34^. While these studies establish important information about cell behavior, site dynamics, and the factor composition of the niche, a functional mechanism pertaining to the inflammation at the site has not been investigated. These observations motivated the development of our model as a hypothesis-testing space for our questions pertaining to inflammation at the site^20^. To create an interpretable space that best reflects the unique niche of recurrent tissue, the RAMI model was built and calibrated on experimental data from our previous work involving macrophage dynamics and their relationship to tumor recurrence^5,6^. Ziraldo et al. established inflammation and wound healing as dynamic systems in which feedback among inflammatory processes can generate divergent outcomes^19^, while more recent ABMs have explicitly demonstrated that altered inflammatory resolution or prolonged inflammatory signaling can shift tissue toward chronic wound or fibrotic states^19,20,28,35^.

### Transient cellular behavior can become persistent tissue state

Here, we find that the tissue’s ability or lack thereof to resolve inflammation is key to creating the recurrent ecology of the site. In testing alternative tissue-memory formulations, we demonstrated that neither indefinite retention nor exclusively recent recollection of inflammation captured biologically-relevant accumulation and resolution of the site^1,6,8^. Previous findings suggest that one key difference between the nonrecurrent and recurrent site begins at the point which wound healing should begin to decrease inflammation and allow the damage to resolve^3,5,6,9^. If the memory used does not offer the tissue the opportunity to resolve, then the model could never reflect a scenario where tissue inflammation is controlled. Instead, the model required that the tissue is given the ability to accrue inflammation, but also progressively resolve it. Due to this separation, we should consider not only tissue memory but also ability to resolve as distinct components of recurrence^1,19,20^. This distinction is consistent with wound-healing models in which the timing and persistence of inflammatory signals alter long-term tissue outcome. Recent work by Chandrasegaran *et al.* demonstrated that the trajectory of the wound healing process can be significantly altered simply by varying the timing of senescent cell signaling^19^. Similarly to their use of inflammatory senescence-associated secretory phenotype (SASP) as a representative for more granular SASP-associated signaling such as mixtures of cytokines and growth factors^36–38^, RAMI extends this general principle by encoding the accumulated consequences of inflammatory remodeling as a latent tissue state.

### Infiltrating cells receive dynamic signals from altered tissue

We developed RAMI to be both receiver and repository of inflammatory tissue memory, creating a tissue-state intermediary that integrates memory in the tissue to elicit changes which impact macrophage phenotype and tumor colonization. This is achieved in part by IL6^6,10^ as an environmental factor emerging from the tissue. Previous tumor-immune models have demonstrated how microenvironmental shifts impact macrophage plasticity and how resultant tumor-macrophage or stromal-macrophage interactions generate emergent tissue behaviors^18,27,39–42^. Similarly, ABMs of inflammatory injury and fibrosis have incorporated feedback between immune-cell behavior and altered tissue state^19,35,39^. The cells changed the tissue, and it was then necessary for the altered tissue to change the cells to complete the mechanism^13^. RAMI provides a computational intermediary between transient cellular events and long-term tissue evolution.

Our ablation experiments distinguish this feedback mechanism from relationships imposed by model construction. RAMI-to-IL6 signaling was explicitly encoded, so increased IL6 should not itself be interpreted as an independent model prediction. In contrast, the population-level macrophage response emerged from stochastic phenotype transitions within the coupled system. Preventing RAMI-derived IL6 from influencing macrophage phenotype significantly reduced the M2/M1 ratio despite continued accumulation of RAMI and generation of IL6. Moreover, feedback ablation reduced subsequent RAMI accumulation itself, demonstrating that tissue-to-cell signaling reinforced later inflammatory remodeling. Thus, the recurrent macrophage ecology depended not only on accumulation of inflammatory memory, but also on closing the loop through which the altered tissue influenced subsequently infiltrating cells.

The sensitivity and robustness analyses further clarify what can and cannot be inferred from the RAMI parameterization. The exploratory sensitivity screen showed that model behavior remains parameter-sensitive, so the hypothesis-driven coefficients should not be interpreted as interchangeable or biologically identified constants. However, the higher-replicate confirmation showed that substantially different combinations of remodeling, persistence, and IL6-feedback parameters can reproducibly generate similar recurrence-associated macrophage states. This indicates equifinality within the model: the experimentally observed macrophage phenotype does not uniquely identify a single underlying parameterization. Accordingly, the present model provides stronger support for a mechanistic class of inflammatory-memory feedback than for the specific numerical values assigned to α, β, persistence, IL6 gain, or IL6 coupling. This distinction is important because these parameters represent hypothesized strengths of biological processes and not directly measured quantities. Additional experimental constraints on inflammatory resolution, macrophage remodeling, and tissue-to-cell signaling will be required to distinguish parameterizations that generate similar macrophage phenotypes.

Our understanding of the site evolved to consider transient immune behavior as the catalyst to a persistent tissue state which then impacts future immune behavior, altering the fate of the site and determining the state. Existing wound-healing models have represented persistent inflammatory mediators, tissue damage, ECM remodeling, and secretory states, while tumor-immune ABMs commonly represent dynamic interactions among macrophages, lymphocytes, stromal cells, and tumor cells^14,18,19,27,28,35,39^. RAMI differs in that it does not represent a specific molecular species or cellular population. Instead, it represents the accumulated consequences of previous inflammatory remodeling and carries that information forward to influence subsequent tissue behavior. To our knowledge, this is among the first computational frameworks to explicitly represent inflammatory tissue memory as a latent state variable linking transient immune behavior to long-term tissue evolution and eventual tumor permissiveness and infiltration.

### Biological Implications

The present work indicates that recurrence may arise not from persistent activation of immune cells alone but from the persistent reprogramming of the tissue they inhabit, supporting previous findings^7,8^. Once inflammatory information becomes embedded within the tissue microenvironment, subsequent immune cells encounter an environment that has already been reprogrammed, creating a self-reinforcing inflammatory niche. The ablation results support that persistent inflammatory burden alone may not be sufficient to maintain this state. Recurrence-associated ecology requires accumulated tissue remodeling to remain biologically accessible to subsequently recruited immune cells. This distinction suggests that interventions capable of disrupting tissue-to-immune feedback may alter the recurrent niche even after inflammatory remodeling has accumulated.

### Limitations and Future Directions

Modeling the recurrent niche was necessary not as a measure to replace experiments but to provide a hypothesis testing framework to complement our ongoing work. The observations here cannot be directly observed in a single experiment. Such work has both a high demand on time and resources. Utilizing computational tools decreases this burden, leading to faster innovation. Despite this, there are hurdles which must be overcome so that the tool can be as useful as possible. Reducing the tissue to the bare agents and functions is often done so that mechanisms within the system can be clearly observed in order to ease parameterization, to be able to interpret the outcomes of model values, and to decrease computational burden^15,16,18,19^. However, critical biological components, such as fibroblasts, the ECM, and the variety of cytokines, are not directly represented. To that end, the model was built with flexibility in mind so that future studies can easily incorporate additional factors and cells. A 2D representation was selected because the available experimental data supported tissue-level rather than fully spatial reconstruction. Our data was sufficient to create the model, and the model was built around the data which roots it in knowns. However, we were limited by the available temporal data. In the future, the model should be expanded to include stromal cells (e.g. fibroblasts, adipocytes), ECM, chemokines, and vascular biology variables such as hypoxia and angiogenesis^7,42–47^. This would allow us to test intervention-style scenarios such as exogenous IFNγ-mediated macrophage reprogramming with higher biological realism^48–51^. As the model evolves, future iterations will be paired with patient-specific data to simulate recurrence and determine potential patient interventions.

The available experimental data cannot uniquely constrain the hypothesis-driven coefficients governing RAMI accumulation and feedback. Sensitivity analysis identified multiple parameter combinations capable of generating similar recurrent macrophage phenotypes, indicating parameter non-identifiability within the current experimental constraints. Accordingly, the present model supports the proposed feedback architecture rather than unique numerical estimates of inflammatory persistence or coupling strength. Although simplified, the model establishes a flexible computational framework that can readily incorporate additional biological variables, which will save time and resources for wet lab experiments. This will lead to innovation and rapid hypothesis testing to reach patient-level interventions to decrease the rate of recurrence for high-risk, immunodeficient TNBC patients. More broadly, this work demonstrates that computational models can reveal mechanisms that are difficult to observe experimentally by integrating diverse biological observations into a single mechanistic framework. Our model indicates that inflammatory tissue memory may represent the missing intermediary between transient immune remodeling to persistent recurrence permissiveness, providing a means for studying radiotherapy-associated recurrence and generating experimentally testable hypotheses for future investigation.

## Conclusion

We establish the latent state variable RAMI as a novel approach to representing unresolved inflammation at tissue sites. Our RAMI model supports a mechanistic framework of recurrence in the context of TNBC. Following radiotherapy, transient immune modeling events are sufficient to cause embedded, long-term alterations to the tissue as inflammatory tissue memory. The state switches from nonrecurrent to recurrent due to additional factors beyond macrophage abundance and phenotype balance that had been identified in previous experimental models. The pro-recurrent niche emerges as a function of immune cell dynamics, impaired tissue inflammation resolution, and crosstalk from the tissue, which create a feedback loop that reinforces recurrence-associated characteristics.

## Acknowledgements

We thank the Translational Pathology Shared Resource core facility for *ex vivo* sample preparation (P30CA068485). The schematics were prepared using BioRender, the NIH BioArt resource, and the noun project, and Canva Free Content library icons.

## Data Availability

The datasets used and/or analyzed for the present work are available from the corresponding author on reasonable request.

---

**Supplementary Table S1.** Parameters and Parameter Sources.

| Parameter Name |  |  |  |  |  |
| --- | --- | --- | --- | --- | --- |
| Macrophage Count | Time Post-Radiation (Days) | Nonrecurrent (Count per Field of View) | $\frac{\text{Nonrecurrent}(t)}{\text{Nonrecurrent}(0)}$ | Recurrent (Count per Field of View) | $\frac{\text{Recurrent}(t)}{\text{Nonrecurrent}(0)}$ |
| Parameter Description | 0 | 5.00 | 1.00 | 3.50 | 0.700 |
| F4/80+ macrophages found in the immunodeficient irradiated mammary fat pad of mice via immunohistochemistry | 1 | 6.41 | 1.28 | 4.43 | 0.866 |
| Source | 5 | 7.59 | 1.52 | 7.08 | 1.416 |
| Rafat *et al* 2018, Cancer Research<br>PMID: 29880480 | 10 | 17.66 | 3.53 | 36.19 | 7.238 |
| Parameter Name |  |  |  |  |  |
| Percent Macrophage Polarization | Family |  |  | Nonrecurrent (%) | Recurrent (%) |
| Parameter Description | F4/80+ Macrophages>M0 |  |  | 90 | 85 |
| Polarization of macrophages at the irradiated mammary fat pad 10 days post-radiation via flow cytometry | F4/80+ Macrophages>M1 |  |  | 5 | 10 |
| Source | F4/80+ Macrophages>M2 |  |  | 5 | 5 |
| Hacker *et al* 2023, Cell Mol Bioengineering<br>PMID: 37810999 |  |  |  |  |  |

